# Community-based Sero-prevalence of Hepatitis B and C Infections in South Omo Zone, Southern Ethiopia

**DOI:** 10.1101/533968

**Authors:** Adugna Endale, Woldearegay Erku, Girmay Medhin, Nega Berhe, Mengistu Legesse

**Affiliations:** Aklilu Lemma Institute of Pathobiology, Addis Ababa University, Addis Ababa, Ethiopia; School of Medicine, College of Medicine and Health Sciences, Dire Dawa University, Dire Dawa, Ethiopia; Department of Microbiology, Immunology and Parasitology, School of Medicine, College of Health Sciences, Tikur Anbessa Hospital, Addis Ababa University, Addis Ababa, Ethiopia

**Keywords:** community, sero-prevalence, hepatitis, Southern Ethiopia

## Abstract

**Background:** Hepatitis B Virus (HBV) and Hepatitis C Virus (HCV) are the leading causes of liver-related morbidity and mortality throughout the world. The magnitude of HBV and HCV infections in Ethiopia has not been well studied at community level. This study aimed at investigating the sero-prevalence and associated risk factors of HBV and HCV among HBV unvaccinated community members in South Omo Zone, Southern Ethiopia.

**Methods:** A community-based cross-sectional study was conducted in three districts from March to May 2018. Structured questionnaire was used to collect relevant clinical and socio-demographic data. Three milliliter of blood sample was collected from each study participant and screened for HBV and HCV using one step hepatitis B surface antigen (HBsAg) test strip and one step HCV test strip, respectively. Samples found positive for HBsAg were further tested using immunoassay of Alere Determine^TM^ HBsAg (Alere Inc., USA). Data was analyzed using SPSS version 25.0.

**Results:** A total of 625 (51.4% males, age 6-80 years, mean age ± SD = 30.83 ± 13.51 years) individuals participated in the study. The sero-prevalence for HBV infection was 8.0% as detected using one step HBsAg test strip, while it was 7.2% using Alere Determine^TM^ HBsAg test. The sero-prevalence for HCV infection was 1.9%. Two (0.3%) of the participants were seropositive for both HBV and HCV infections. High sero-prevalence for HBV infection was associated with weakness and fatigue (AOR = 5.20; 95% CI: 1.58, 17.15), while high sero-prevalence of HCV infection was associated with age group between 46 and 65 years (AOR = 13.02; 95% CI: 1.11, 152.41).

**Conclusion:** this study revealed higher-intermediate endemicity level of HBV infection and low to intermediate endemicity level of HCV infection in the study area. Clinical symptoms like weakness and fatigue were found to be indictors for HBV infection, while individuals in the age group between 46 and 65 years were at higher risk for HCV infection. Provision of community-based health education, vaccination, mass screening and providing treatment would have utmost importance in reducing the transmission of these diseases in the present study area.

## Introduction

Hepatitis, inflammation of the liver, can be caused due to infectious and non-infectious agents such as viruses, bacteria, fungi, parasites, alcohol, drugs, autoimmune diseases, and metabolic diseases [1]. The most common causes of hepatitis are viruses; namely hepatitis A, B, C, D and E viruses. Among these, hepatitis B virus (HBV) and hepatitis C virus (HCV) are the most common causes of viral hepatitis [2].

Both HBV and HCV can be transmitted through sexual, blood or vertically from mother-to-child [3]. Thus, individuals who require blood transfusion, those who have multiple sexual partners and infants born to HBV or HCV infected mothers are at a high-risk for acquiring HBV or HCV infection [4]. Both viruses can cause acute and chronic infection of the liver [5, 6]. Chronic HBV and HCV infections are the leading causes of liver-related morbidity and mortality [7, 8]. Between 15% and 40% of chronically infected individuals can develop serious liver disease and transmit the viruses to others [9, 10]. Globally, about 257 million people were living with HBV infection, and 71 million people were living with HCV infection in 2015 [4, 11]. About 1 million people die each year from cases related to viral hepatitis [4]. An estimated 50% to 80% of cases of primary liver cancer result from infection with HBV [12, 13].

A safe and effective vaccine for HBV has been available since 1982, whereas no vaccine exists for HCV [14]. Treatment options for both viruses are advancing rapidly, and several new antiviral drugs have become available [15]. By the end of 2015, only 9% of HBV-infected people and 20% of HCV-infected people had been diagnosed. Of those 1.7 million people who found positive for HBV infection, only 8% were treated, while only 7% were treated among 1.1 million people who were positive for HCV infection in 2015 [4]. The global targets for 2030 are to diagnose 90% of people with HBV and HCV infections and treat 80% of eligible patients [16].

In Ethiopia, more than 60% of chronic liver disease and up to 80% of hepatocellular carcinoma (HCC) are due to chronic HBV and HCV infections [17]. According to WHO, Ethiopia is among hepatitis endemic countries in the world with intermediate to hyperendemic endemicity level [18]. However; Ethiopia is regarded as a country with no national strategy for surveillance, prevention and control of viral hepatitis. Above all Ethiopian children including children in our current study site have not had access to vaccination against HBV. Data on the epidemiology of HBV and HCV infections in Ethiopia at the community level are scarce. Particularly, HCV infection remains under-diagnosed and under-reported, despite its high infectious nature. Thus lack of adequate epidemiological data at the community level on hepatitis in Ethiopia can affect the global targets to control HBV and HCV infections. For this reason, assessing the sero-prevalence of HBV and HCV infections at community level is very important to develop strategies to reduce transmission among the community members. Therefore, this study was aimed at investigating the sero-prevalence and associated factors for HBV and HCV infections among HBV unvaccinated community members in South Omo Zone, Southern Ethiopia.

## Materials and Methods

### Study design, period and setting

A community-based cross-sectional study was conducted in three districts (Hamer, Debub Ari and Bena-Tsemay) of South Omo Zone, Southern Ethiopia from March to May 2018. The study area is located about 750 Kilometers from Addis Ababa, the capital city of Ethiopia. According to the 2007 Ethiopian census, the total population of South Omo Zone is 647,655 [19] and the eight largest ethnic groups are Ari (44.59%), Male (13.63%), Dasenech (8.17%), Hamer (8.01%), Bena (4.42%), Amhara (4.21%), Tsemai (3.39%), and Nyangatom (2.95%) [20]. Most of the inhabitants are nomadic pastoralists in the Hamer District and farmers in the Debub Ari and Bena-Tsemay districts [19]. The study districts were purposely selected because of their endemicity for yellow fever outbreaks [21], which has similar clinical features with viral hepatitis.

### Study participants, sample size estimation and sampling method

The study participants were individual’s age 5 years and above, residents of the study districts and volunteered to participate in the study. Sample size for the nomadic population (Hamer District) was calculated with the estimated HBsAg sero-prevalence of 7.4 [22] with 95% confidence, 5% margin of error, 15% non-response rate and 1.5 design effect. For the mixed farming and settled farming population (Debub Ari and Bena-Tsemay Districts) the sample size was calculated with the assumptions: sero-prevalence of HBsAg in the community is 10.5% [22] at confidence level of 95% and 5% margin of error; estimated non-response rate 10% and design effect of 2.0. Accordingly, these resulted in minimum sample size of 621. Based on these estimation 306 individuals from Hamer District and 319 from Debub Ari and Bena-Tsemay districts were involved. The households from each district were selected using systematic random sampling technique after getting the n^th^ value by dividing the total number of households by the sample size. From each selected households one participant was randomly recruited using lottery method.

### Data collection

Three milliliter of blood sample was collected from each of the study participants, and serum was separated and stored at appropriate temperature until screened for HBV and HCV infections. Structured questionnaire was administered to collect socio-demographic and relevant clinical data. One step HBsAg test strip (Nantong Diagnos Technology Co., Ltd., China) was used for the screening hepatitis B surface antigen (HBsAg). Whereas, one step HCV test strip (Nantong Diagnos Technology Co., Ltd., China) was used to detects antibodies against HCV following the instructions of the manufacturer. Samples found positive for HBV infection by one step HBsAg test strip were further screened using qualitative immunoassay Alere Determine^TM^ HBsAg (Alere Inc., Massachusetts, USA) which is more specific but less sensitive than the One Step HBsAg test [23–25]

### Data entry and analysis

Data was entered using Epi-Data Entry version 3.1 and analyzed using SPSS version 25.0. Descriptive statistics; mean and standard deviation for continuous variables and frequency for categorical variables were used. Bivariable and multivariable logistic regression analysis were used to assess factors associated with sero-prevalence for HBV and HCV infections. Variables which showed association in multivariable analysis were considered as final predictors of the dependent variable. All tests were performed at 5% level of significance.

### Ethical considerations

The study was carried out after getting ethical approval from the Institutional Review Board (IRB) of Aklilu Lemma Institute of Pathobiology, Addis Ababa University. Then, data was collected after getting permission from South Omo Zone and district health offices. The objective, the possible risks & benefits of the study were explained to the participants or their guardians in local languages and informed written consent was obtained from the participants or from their guardians. The participants were assured that they had full right to participate or not to participate in the study. Sero-positive individuals were advised and linked to health facilities to obtain appropriate treatment and care. All information obtained in this study was kept confidential.

## Results

### Socio-demographic characteristics of the study participants

A total of 625 participants (51.4% males, age range from 6 to 80 and mean age ± SD = 30.83 ± 13.51 years) participated in this study. Table 1 shows the socio-demographic characteristics of the study participants.

**Table 1:**
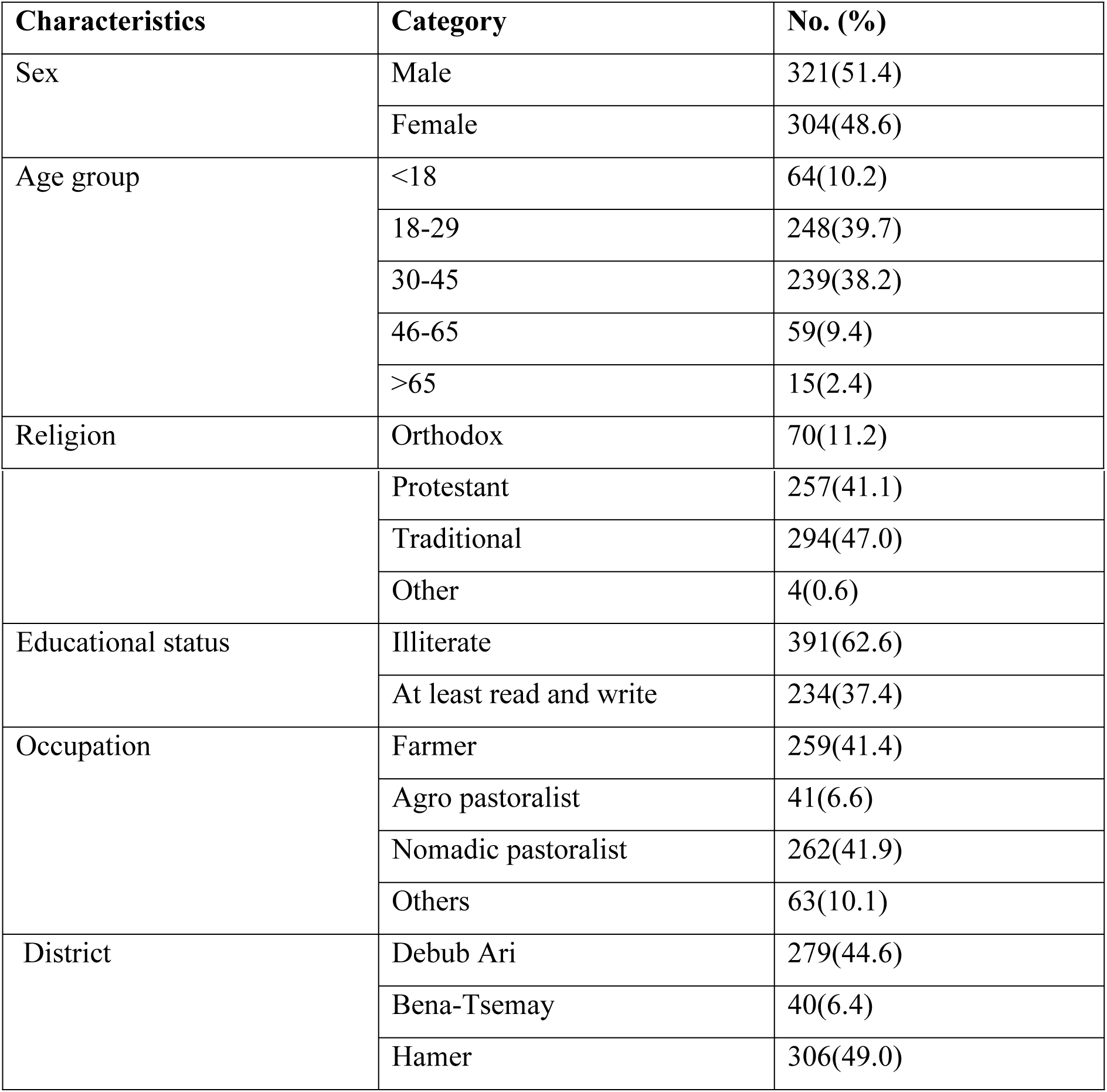
Socio-demographic characteristics of the study participants

### Clinical sign and symptoms reported by the study participants

As shown in table 2, the most common sign and symptoms reported were upper abdominal pain especially on the right side (18.9%) followed by joint pain and muscle aches (16.0%) and weakness and fatigue (7.5%).

**Table 2:** Sign and symptoms reported by the study participants

| Characteristics | Category | No (%) |
| --- | --- | --- |
| Upper abdominal pain | Yes | 118(18.9) |
|  | No | 507(81.1) |
| Dark urine | Yes | 43(6.9) |
|  | No | 582(93.1) |
| Joint pain and muscle aches | Yes | 100(16.0) |
|  | No | 525(84.0) |
| Loss of appetite | Yes | 17(2.7) |
|  | No | 608(97.3) |
| Nausea and vomiting | Yes | 26(4.2) |
|  | No | 599(95.8) |
| Weakness and fatigue | Yes | 47(7.5) |
|  | No | 578(92.5) |
| Yellowing of skin and eyes (jaundice) | Yes | 11(1.8) |
|  | No | 614(98.2) |
| Other non-specific symptoms such as fever, headache and body rash | Yes | 14(2.2) |
|  | No | 611(97.8) |

### Sero-prevalence for HBV and HCV infections

The overall sero-prevalence for HBV infection was 8.0% (9.4% among males and 6.6% among females) using one step HBsAg test strip and 7.2% (8.8% among males and 5.6% among females category) using HBsAg Alere Determine^TM^ test. Among 50 samples found positive for HBV infection by one step HBsAg test strip, 5 samples were found negative by HBsAg Alere Determine^TM^ test. The overall sero-prevalence for HCV infection was 1.9% (2.2% among males and 1.6% among females). Two (0.3%) of the participants were co-infected with both HBV and HCV. The sero-prevalence of HBsAg and anti-HCV was relatively higher in Bena-Tsemay district. Under the bivariable analysis, HBV infection was significantly associated with weakness and fatigue (P<0.01) and HCV infection associated with age group (P=0.02) (Table 3).

**Table 3:**
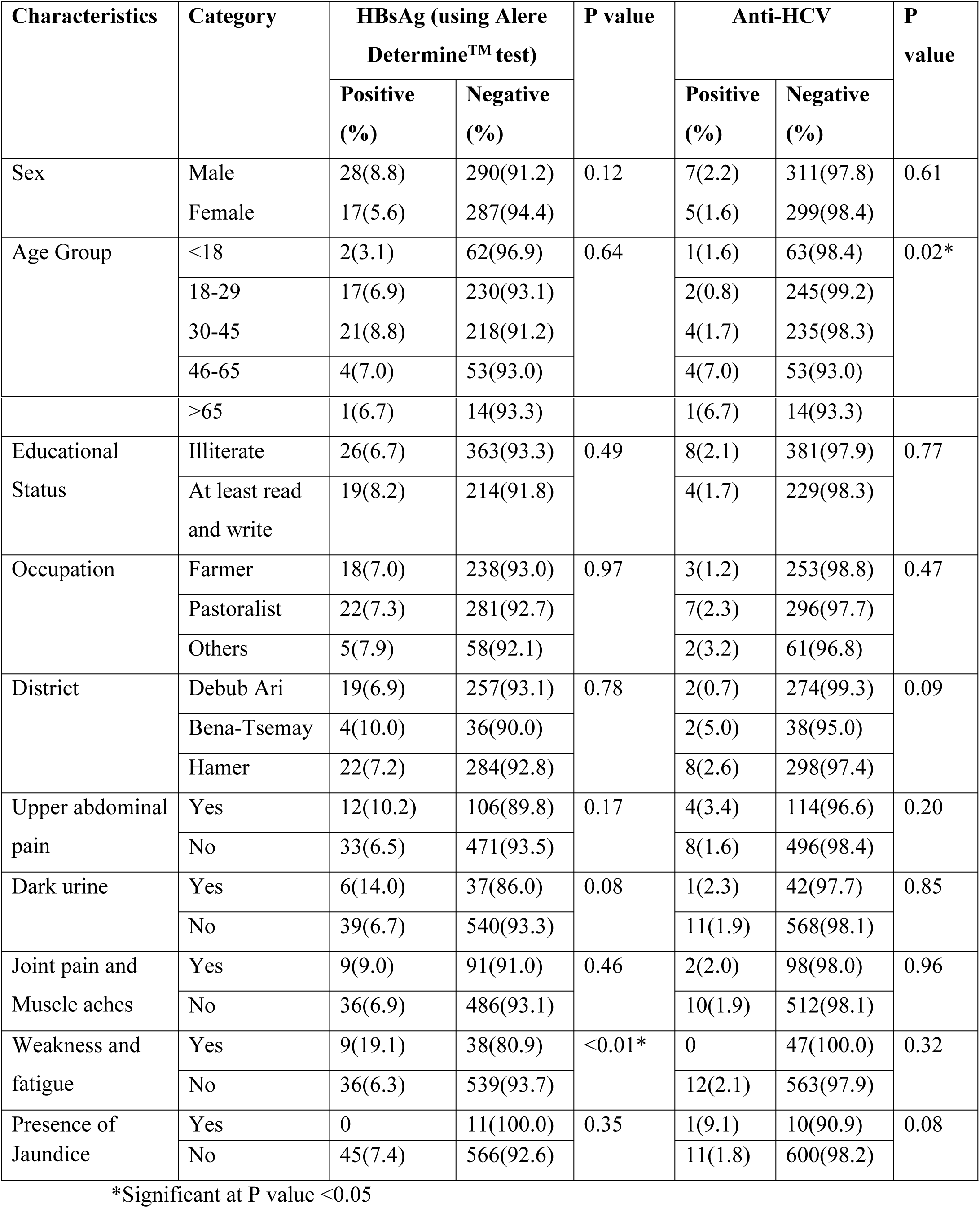
Sero-prevalence for HBV and HCV infections

### Independent Predictors of HBV and HCV Infections

A multivariable logistic regression analyses was performed to explore the independent predictors of HBV and HCV infections. Having body weakness and fatigue (AOR = 5.20; 95% CI: 1.58, 17.15) and age group 46-65 (AOR = 13.02; 95% CI: 1.11, 152.41) were significantly associated with HBV (Table 4) and HCV infections respectively at P value <0.05 (Table 5).

**Table 4:** Independent predictors of HBV infection

| Characteristics | Category | HBsAg |  |
| --- | --- | --- | --- |
|  |  | COR (95% CI) | AOR (95% CI) |
| Sex | Male | 1.63(0.87, 3.04) | 1.45(0.75, 2.79) |
|  | Female | Reference | Reference |
| Age Group | <18 | Reference | Reference |
|  | 18-29 | 2.29(0.52, 10.19) | 1.51(0.31, 7.43) |
|  | 30-45 | 2.99(0.68, 13.09) | 2.10(0.42, 10.49) |
|  | 46-65 | 2.34(0.41, 13.28) | 1.92(0.30, 12.50) |
|  | >65 | 2.21(0.19, 26.17) | 1.52(0.12, 20.07) |
| Educational Status | Illiterate | 0.81(0.44, 1.49) | 0.79(0.38, 1.68) |
|  | At least read and write | Reference | Reference |
| Occupation | Farmer | Reference | Reference |
|  | Pastoralist | 1.04(0.54, 1.98) | 1.01(0.31, 3.30) |
|  | Others | 1.14(0.41, 3.20) | 1.33?(0.3, 5.08) |
| District | Debub Ari | Reference | Reference |
|  | Bena-Tsemay | 1.50(0.48, 4.67) | 1.24(0.33, 4.76) |
|  | Hamer | 1.43(0.47, 4.40) | 1.87(0.45, 7.78) |
| Upper abdominal pain | Yes | 1.62(0.8, 13.23) | 1.63(0.62, 4.27) |
|  | No | Reference | Reference |
| Dark urine | Yes | 2.25(0.89, 5.64) | 1.51(0.44, 5.20) |
|  | No | Reference | Reference |
| Joint pain and | Yes | 1.34(0.62, 2.87) | 2.03(0.58, 7.05) |
| **Muscle aches** | No | Reference | Reference |
| **Weakness and fatigue** | Yes | 3.55(1.59, 7.900 | 5.20(1.58, 17.15)* |
|  | No | Reference | Reference |

CI (confidence interval), COR (crude odds ratio), AOR (adjusted odds ratio) and \*(significant at p<0.05)

**Table 5:** Independent predictors of HCV infection

| Characteristics | Category | Anti-HCV |  |
| --- | --- | --- | --- |
|  |  | COR (95% CI) | AOR (95% CI) |
| **Sex** | Male | 1.35(0.42, 4.29) | 1.15(0.33,4.02) |
|  | Female | Reference | Reference |
| **Age Group** | <18 | Reference | Reference |
|  | 18-29 | 0.51(0.05, 5.76) | 0.59(0.05, 7.48) |
|  | 30-45 | 1.07(0.12, 9.76) | 2.14(0.20, 23.28) |
|  | 46-65 | 4.76(0.52, 43.85) | 13.02(1.11, 152.41)* |
|  | >65 | 4.50(0.27, 76.38) | 4.62(0.16, 137.01) |
| **Educational Status** | Illiterate | 1.20(0.3, 64.04) | 1.59(0.34,7.49) |
|  | At least read and write | Reference | Reference |
| **Occupation** | Farmer | Reference | Reference |
|  | Pastoralist | 1.99(0.51, 7.79) | 3.81(0.24, 60.11) |
|  | Others | 2.77(0.45, 16.91) | 1.36(0.05, 36.52) |
| **District** | Debub Ari | Reference | Reference |
|  | Bena-Tsemay | 7.21(2.00, 52.70) | 11.20(1.07, 117.27) |
|  | Hamer | 1.96(0.40, 9.57) | 18.20(0.65, 513.25) |
| Upper abdominal pain | Yes | 2.18(0.64, 7.35) | 1.67(0.36, 7.72) |
|  | No | Reference | Reference |
| Dark urine | Yes | 1.23(0.16, 9.75) | 1.47(0.13,16.22) |
|  | No | Reference | Reference |
| Joint pain and Muscle aches | Yes | 1.05(0.23, 4.84) | 2.26(0.37,13.69) |
|  | No | Reference | Reference |
| Presence of Jaundice | Yes | 5.46(0.64,46.38) | 3.50(0.24, 52.29) |
|  | No | Reference | Reference |

CI (confidence interval), COR (crude odds ratio), AOR (adjusted odds ratio) and \*(significant at p<0.05)

## Discussion

This study revealed higher-intermediate endemicity level [26, 27] of HBV infection with overall sero-prevalence of 7.2% among the general population in South Omo Zone. The sero-prevalence of HBsAg is almost similar with the previous national pooled prevalence of 7.4% of Ethiopia [22] and within the range of a prevalence of 5–10% reported among adult population in sub-Saharan African countries [28]. However the prevalence is relatively higher than 3.1% prevalence reported from a recent community-based study done in Gojjam, Northwest Ethiopia [29]; 6.1% the whole African region and 3.5% global prevalence among the general population [4]. Thus the relative increase in the prevalence of HBsAg indicates that the area should be one among the priority target areas for the prevention and control of hepatitis.

In the case of HCV sero-prevalence, anti-HCV detection rate varied from low to higher– intermediate endemicity levels [26, 27] among the different districts with the highest sero-prevalence (5.0%) detected in Bena-Tsemay district. The overall sero-prevalence of anti-HCV in the study area was 1.9%. The overall sero-prevalence of anti-HCV recorded in our study area was less than that from the pooled national prevalence of 3.1% [22] and 3.0% reported in Sub-Saharan Africa [30]. However, it is still greater than the 1.0% prevalence reported from Gojjam, Ethiopia [29]; 0.3% in Djibouti, 0.9% in Somalia, and 1.0% in Sudan [31] among the general populations. Although the overall sero-prevalence of HCV infection in the study area is considered to be low according to the WHO classification [26, 27], relatively higher prevalence detected in Bena-Tsemay district indicates it is a marked public health problem in that district.

Since HBV and HCV share common route of transmission, co-infection between the two viruses is common, especially in high endemic areas and among people at high-risk of parenteral infection [32]. In this study co-infection between HBV and HCV infection serologic markers was 0.3% among the general population. This is in contrast to some studies from Ethiopia where no serologic markers for HBV and HCV co-infection was reported [33-35], indicating that co-infection between HBV and HCV may not be uncommon in the areas where both HBV and HCV are endemic. Our finding is supported by findings from many other studies conducted elsewhere which reported dual infection of HBV and HCV: 0.7% in Nigeria among prison inmates [36]; 5.9% and 2.0% in India among patients with chronic liver disease [37] and hemodialysis patients [32] respectively and 7.7% in Mongolia among patients with chronic liver disease [38].

Under the multivariate analysis, having body weakness and fatigue was independently associated with HBV serological marker (HBsAg) exposure status. Those participants having body weakness and fatigue were almost five times at higher risk of being positive for HBsAg as compared to their counter parts without such problem. Previously numerous studies revealed that fatigue (generalized body weakness) is the most commonly reported symptom in patients with viral hepatitis [39-42]. Worth noting here, however, since a good number of clinical symptoms of viral hepatitis overlap with those of arboviral infections [43], involvement of the later cannot be overruled. In fact, the study area is among the known hot spots for yellow fever outbreaks [21] and is also considered to be an endemic site for many arboviral diseases because of its proximity to neighboring endemic/enzootic country Kenya, where repeated arboviral disease outbreaks have been reported [44]. Moreover, multivariate analysis also indicated that those of participants in the age group 46 to 65 years were more than thirteen times at higher risk of being positive for HCV infection as compared to those in the age group less than 18 years. These implying, older individuals are at higher risk of getting HCV infection as compared to younger. The observed significantly higher sero-prevalence in older individuals might be attributed to frequency of exposure. Similar findings were reported from studies done in: Rwanda [45], Egypt [46] and Madagascar [47].

### Limitations

Our serologic assays for HBV and HCV detection did not provide evidence of active viremia and identification of infected individuals in the antibody-negative phase (those in the window period) for HCV detection.

### Conclusion

In general, the finding in this study revealed a higher-intermediate HBV and a low to intermediate HCV infection endemicity levels among the general population in South Omo Zone. Those individuals having body weakness and fatigue and older individuals (age group 46-65) were at higher risk of acquiring HBV and HCV infections, respectively. These observations call for responsible health policy makers to develop a practical plan of intervention with the goal of prevention and control of these infections, such as screening those belonging to the high-risk group among the general population, improvement in the expansion of HBV vaccination, and provision of health education. Further research using molecular and other more sensitive and specific assays for detecting active HBV and HCV infections is also needed for the future.

## Acknowledgements

The authors would like to acknowledge the study participants and South Omo Region and Districts Health Administration.

## Author Contributions

### Conceptualization and design

Adugna Endale, Mengistu Legesse, Woldearegay Erku, Girmay Medhin, Nega Berhe.

### Acquisition of data

Adugna Endale, Mengistu Legesse, Woldearegay Erku, Girmay Medhin, Nega Berhe.

### Formal analysis and interpretation

Adugna Endale, Mengistu Legesse, Woldearegay Erku, Girmay Medhin, Nega Berhe.

### Writing - original draft

Adugna Endale.

### Writing - review & editing

Adugna Endale, Mengistu Legesse, Woldearegay Erku, Girmay Medhin, Nega Berhe.

